# A training protocol for human classification of Asian elephant images from trail cameras

**DOI:** 10.64898/2026.09.24.754213

**Authors:** Finley Daecher, Sateesh Venkatesh, Carolina Loera, Alyssa Ayan, U.S. Weerathunga, T.V. Pushpakumara, Malaka Abeywardana, U.L. Thaufeek, Shermin de Silva

## Abstract

Trail cameras have become ubiquitous tools for ecological data collection over recent decades. Despite progress in the development of automated algorithms and artificial intelligence for image classification, our ability to process large volumes of data remain limited by the need for trained human observers to make refined judgements. We provide guidance on placement of trail cameras for observing Asian elephants (*Elephas maximus)* and outline a protocol for training and testing naïve human observers in performing image classifications (age/sex class and group composition) that cannot yet be automated. This process can be used to develop a high-throughput workflow capable of extracting useful data from large volumes of images. Our training material consisted of 14,007 images collected from 6 trail cameras around Udawalawe National Park in Sri Lanka from 2017-2019. In the first stage, expert observers (n=3) trained a group of inexperienced participants (n=4), who engaged in an iterative process to develop a protocol document. The document was then tested on a second set of subjects (n=6) each of whom classified 350 test images in four separate sequential batches using quantitative measures of precision and accuracy. The test set was sampled from 54,435 images from an additional 25 cameras. When compared to expert observers, they achieved a fair level of precision (Fleiss’ kappa = 0.247) and 82.6% accuracy. Our approach can usefully be extended to other species and contexts.

## Introduction

Since their commercial introduction in the 1990s, trail cameras have offered a means to conduct non-invasive observations of wildlife across space and time (O’Connell et al., 2011). From 2005 to 2021, studies using trail cameras increased by an annual rate of 1.26, documenting the occurrence and behavior of thousands of species (Delisle et al., 2021). The increased scientific use of camera traps encouraged drastic improvements to image quality and ease-of-use, making them especially valuable for observing cryptic and rare species (McCallum, 2013) and a powerful means of assessing species responses to environmental change (Oliver et al., 2023). In recent years, artificial intelligence (AI) has emerged as a potent tool for camera trap data processing with programs such as Wildlife Insights and Conservation AI (Brickson et al., 2023; Vélez et al., 2023). In particular, there have been major advances in individual-level identification for species with patterned markings (Zábó et al., 2026).

However, AI systems have several limitations. First and foremost, they require large pre-classified training datasets. Second, the required “deep learning” packages can be costly (Green et al., 2020). Mass image identification is also slowed by the simultaneous occurrence of multiple species and images with changing backgrounds (Norouzzadeh et al., 2021a). When compared to Microsoft AI’s MegaDetector software for camera trap data, humans still outperformed the AI in wildlife capture ability and detection size (Leorna & Brinkman, 2022). Finally, although AI has been successfully applied toward tasks such as species recognition and even individual-recognition in a handful of taxa (de Silva et al., 2022; Ma et al., 2025), their feasibility remains limited by the need for training data even when high-quality images are available, especially for non-patterned species (de Silva et al., 2022; but see Norouzzadeh et al., 2021). Moreover, AI has limited capacity for making refined classifications extending beyond the species level for the majority of taxa (e.g. behavior, age-sex classes, or other contextual information). For all these reasons, the need for a human to check the results persists (Vélez et al., 2023). This has led to the increasing use of “citizen science” initiatives, which effectively crowdsource image classifications using untrained or minimally trained observers, and can complement AI (Green et al., 2020; Norouzzadeh et al., 2021b). However, when multiple human participants are involved, inter-observer reliability may be low, requiring systematic effort to evaluate and address (Zett et al., 2022) and sensitive images (e.g. containing human activity) cannot be shared publicly in order to maintain privacy (Sandbrook et al. 2021).

Here we focus on the use of trail cameras to observe Asian elephants (*Elephas maximus*) as our target species. Listed as endangered on the IUCN Red List, improving methods to learn more about the movement and population dynamics of *E. maximus* is increasingly important (Williams et al., 2020). The use of trail cameras can provide insights into elephant movements and behavior in protected areas as well as landscapes of varying disturbance and in human activity (Fernando et al., 2022; Lee et al., 2024; Morrison et al., 2022). They can provide a snapshot of the demographic attributes of populations, such as age and sex-structure as well as characteristics of individual animals engaged in potentially problematic behavior (Ranjeewa et al., 2025; Smit et al., 2019; Varma et al., 2007). Although species-level classification of Asian elephants from camera trap images can be achieved by both AI and naïve human observers with relative ease, individual recognition requires use of 3-dimensional information rather than the use of 2-dimensional patterns (de Silva et al., 2022). Designation of social groupings and age/sex class requires substantial training and can be subjective even for human observers, which impedes automation. Our goals for this study were to (1) pilot placement of trail cameras set up specifically for observing elephants to determine optimal configurations (2) develop a protocol document for inexperienced volunteers to reference for extracting demographic and basic behavioral data from trail camera images of Asian elephants and (3) outline a methodology for developing and testing the document with naïve volunteers that can be applicable for other species and contexts.

## Methods

### Study site and materials

This study focused on the periphery (forest edge) and interior of the Wetahirakanda Sanctuary that connects Udawalawe National Park and Yala National Park block 6 (formerly, Lunugamwehera National Park; Figure 1) in Southern Sri Lanka. The area is surrounded by a matrix consisting primarily of small-holder agricultural lands as well as a few plantations of teak. Small man-made water sources are also located along the periphery, used both by agriculturalists and wildlife. As this pilot study sought to maximize the likelihood of elephant captures rather than to assess elephant occurrence or abundance, cameras were positioned non-randomly along trails and pathways indicated to be used by elephants according to local residents. Whenever cameras were oriented toward agricultural lands with human activity, they were placed with the knowledge and consent of land holders following best practices for responsible data collection (Sandbrook et al. 2021). Cameras were deployed at 6 locations for 34 months between the years 2017-2019, representing a total of 505 trap days.

**Figure 1:**
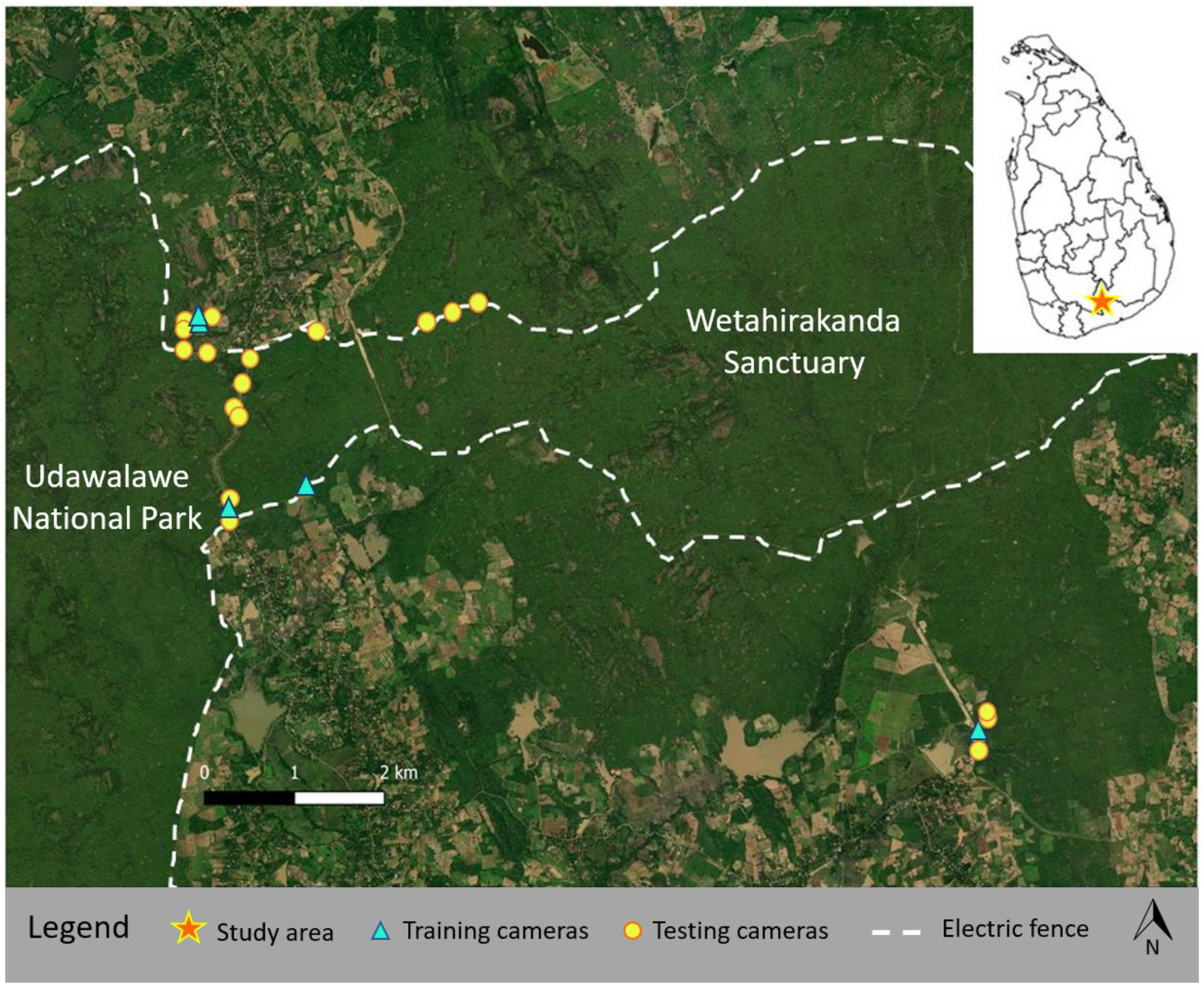
Study area. Wetahirakanda Sanctuary designates the area within the electric fence (dashed line), managed by the Department of Wildlife Conservation. Forest beyond these boundaries are managed by the Forest Department or privately. Cameras (triangles) were placed along the forest edge.

We used Bushnell™ trail cameras (models Trophy Cam HD Essential E2 and E3) housed in protective metal casings. They were powered by rechargeable lithium ion batteries and fitted with 32 GB memory cards. Cameras were attached to trees using cloth straps and locked into place using Python™ cables. Devices were deployed 24 hours a day, activated by motion within 30m during the day and infra-red heat sensors during the nighttime. When activated in low light conditions, the camera uses a red flash that minimizes disturbance to animals. Photos taken using the flash at night show only black and white whereas daytime images show color. Cameras were set to take five 16 mega-pixel photos in rapid succession when activated. The height varied slightly between 1.4-1.6m depending on the terrain and available mounting trees, but was intended to be at the head-level of an elephant to capture distinguishing features, in particular the ears and head shape. During an initial pilot phase of one month, we oriented cameras in 2 different positions relative to the trail: (a) Pairs of cameras were placed directly opposite one another, perpendicular to the trail, with no tilt. Analogous to studies of carnivores, each camera would capture one side of an animal, ensuring both sides were seen. (b) A single camera was placed pointing diagonally across the trail. For the remainder of the study, we used position (b) in all but one location (see results).

### Protocol development

#### Stage 1 – Defining the coding scheme

Experienced observers (SdS, USW, TVP) first developed the image classification categories and protocol. It is necessary to first propose criteria to distinguish among putatively independent events that are captured by cameras. Images were first classified into *event types* (Table 1, supplemental material) and multiple images that represented a sequence from the same event (e.g. a person, animal or group of individuals walking past, see Box 1) were grouped into a unique *event number*. For images of wildlife other than elephants, the species was recorded if it could be identified. If an image contained more than one species, a unique event number was assigned to each species. For images containing elephants, each event was also assigned a *group size* and *group composition* (Table 1). Elephants were differentiated by sex and adult vs. subadult age classes (de Silva et al., 2011b). Younger age classes were not differentiated for purposes of this study as they moved with adults.

**Table 1:** Classification scheme. Species, group size and group type classifications.

| Event type | Group size | Group composition (*E. maximus specific*) |
| --- | --- | --- |
| Elephant | 1 | AM: Solitary adult male |
| Human | 2-5 | AF: Solitary adult female |
| Vehicle | 6-9 | SAM: Solitary subadult male |
| Domestic animals | 10+ | SAF: Solitary subadult female |
| Other wildlife |  | MG: Male group |
|  |  | FG: Female group |
|  |  | XG: Mixed-sex group |
|  |  | AUK: Adult of unknown sex |
|  |  | SAUK: Subadult of unknown sex |
|  |  | UKG: Group of unknown composition |

#### Stage 2 – Training the first set of naïve classifiers

Four volunteers (group 1) with no prior experience observing elephants were trained to make image classifications by experienced observers (SdS and SV) through a collaborative, iterative process with many opportunities to ask for clarifications. Group 1 was then tested on a batch of 100 randomly selected photos, which expert observers had also classified. When they achieved a level of accuracy and precision of >90% (described below), they then collectively classified a subset of 14,007 images from six cameras which included all event types. During this process the students developed a guidance document by themselves, with feedback from the trainers, specifically for the purpose of classifying elephant images (supplementary text 1). This step was critical as it ensured that the features highlighted in the guidance document were not only accurate and valid, but also readily identifiable and could be differentiated by someone with no prior knowledge or experience with the target species.

#### Stage 3 – Validation of guidance document

Discrimination of non-elephant events was straightforward therefore we focused only on validating the guidance developed for elephant images. To test the effectiveness and repeatability of our protocol, we provided the document developed by group 1 to a new set of six volunteers (group 2) with no prior experience. Each individual was then tested with one batch of 50 photos and three subsequent batches of 100 photos in which they were asked to classify event transitions (i.e. change points from one event to the next), group size, and composition by age and sex. The first batch contained a mixture of group compositions from randomly selected photos which did not preserve image sequences for events. Batches 2-4 also contained a mixture of group compositions but preserved image sequences representing intact events. Volunteers in group 2 were not permitted to ask any clarifying questions or collaborate while classifying a batch, but this was done at the end of each successive batch, with concurrent modifications and improvements to the guidance document. Volunteers were asked to self-report the time it took them to complete each batch. We then rated classifications by individuals in group 2 for each batch of photos separately using metrics of *accuracy* and *precision* (Figure 2).

**Figure 2:**
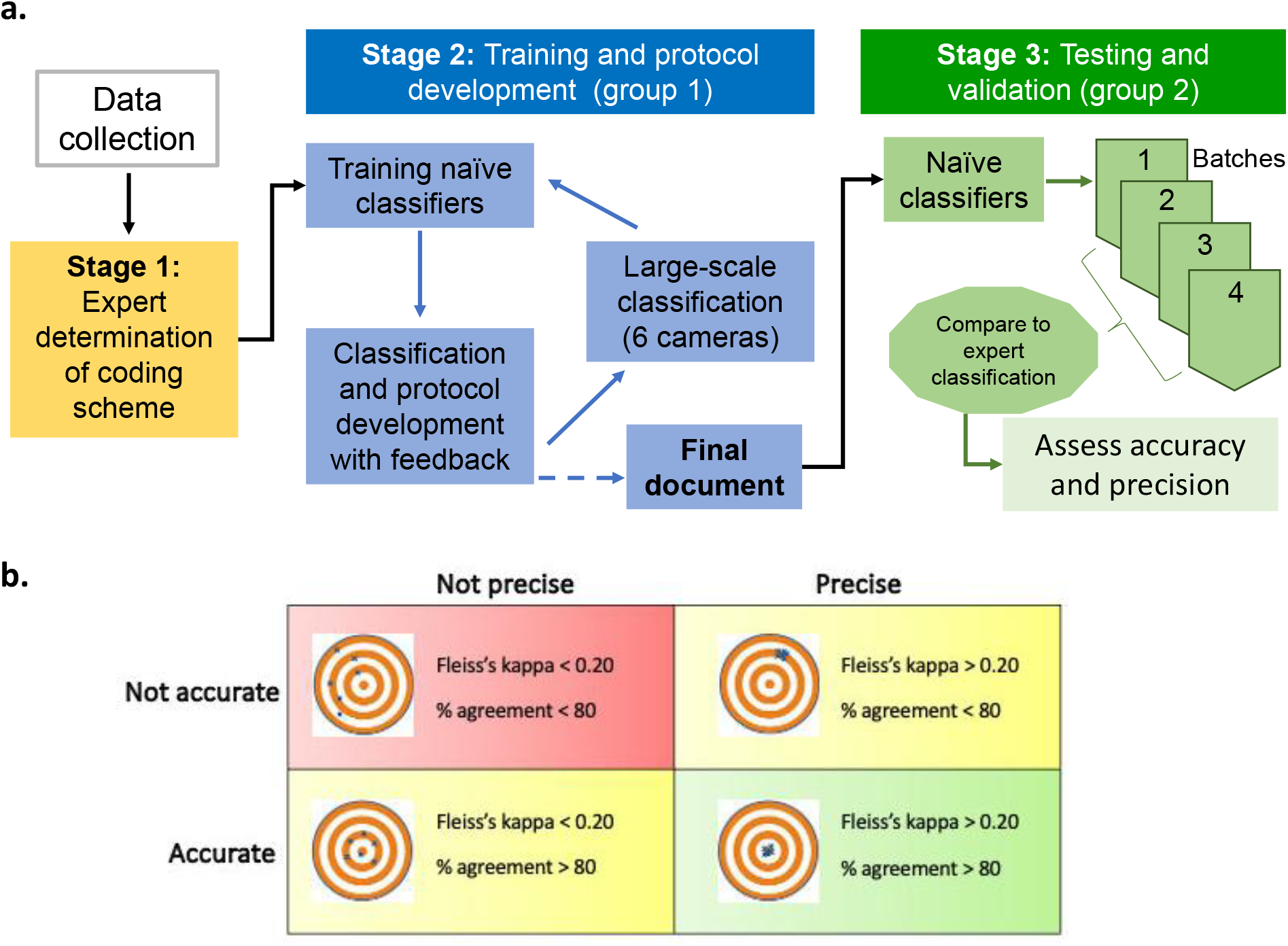
Workflow. a) Training and testing protocol. b) Graphical interpretation of accuracy vs. precision, where the center of the target represents “correct” classifications. This visualization is in itself useful for explaining training goals.

The accuracy of classifications by volunteers was compared to those of experienced classifiers (SdS, SV, TVP). For group types, one point was given for the correct sex (male, female, mixed), and one point was given for the correct age (adult, subadult).

Event changes occur throughout the dataset; one point was given for a correct change in event whereas 1 point was deducted for an incorrect event change (since event changes are variable from one data set to the next, the total points possible changed depending on the batch). One point was given for each correct group size code. One point from the total possible score was deducted for any images in the batch that did not contain elephants or which were ambiguous. Accuracy was determined by dividing the volunteer score by the total points possible (Table 2).

**Table 2:**
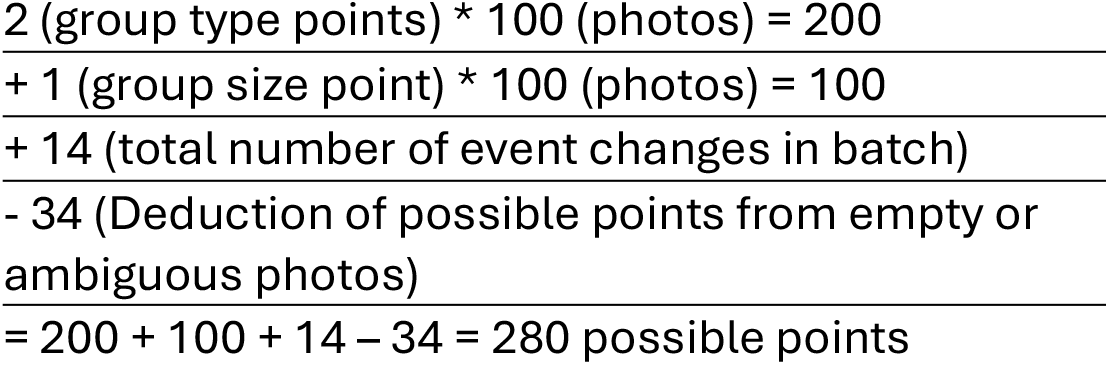
Example calculation of accuracy for a batch of 100 images. In this case, a score of 224 would be needed to achieve 80% accuracy.

To calculate precision, Fleiss’ kappa was used to test if volunteers’ classifications were consistent with one another. Agreement is categorized by kappa values, where values below 0 indicate no agreement, 0-0.20 slight agreement, 0.21-0.40 fair agreement, 0.41-0.60 moderate agreement, 0.61-0.80 substantial agreement, and 0.81-1.00 almost perfect agreement (Nichols et al. 2010). Fleiss’ kappa was calculated using R studio with the package “irr” (Gamer et al. 2019).

## Results

We found that pairing cameras did not work well for several reasons. First, it seemed that having two flashes occurring simultaneously at eye-level on either side of passing elephants caused them to take extra notice of the devices and more likely to attempt to manipulate and dislodge them. Secondly, although in principle it should have been possible to match images of a given individual from one side with images taken from the other, the lack of precise coordination among cameras and the fact that animals were sometimes moving in groups of multiple overlapping individuals made this task extremely impractical even for experienced observers. Lastly, this configuration doubles the number of cameras used per location. The second configuration, in which a camera was pointed diagonally across the trail, was more effective as both ears of the same individual could still sometimes be seen and it allowed camera deployments over more locations. Moreover, placing the cameras slightly above elephant eye-level but with a downward tilt attracted less attention from the target species. We had no cameras lost to theft and none in the pilot sample were damaged by elephants.

Although our target species was Asian elephants, both configurations were equally effective in capturing other non-target species. A total of 5,359 non-target wildlife events were observed. When multiple species occurred in the same image, these were recorded as separate parallel events, with the total being the sum of all events. The other observed species included spotted deer, sambhur deer (rarely sighted at this location), porcupine, leopard, and domestic water buffalo.

### Distinguishing among events

The mean elephant event duration was 2.25 minutes ± 0.21 SE (median: 0.25 minutes), whereas the mean interval between events was 376.67 minutes ± 33.67 SD (median: 110.22 minutes; Figure 3). 83.1% of elephant events (n=2471) fell within a 5-minute time duration, while 90.2% fell within 10 minutes (Figure 3; Note that it is possible for an event to be longer than 10 minutes in duration if multiple successive photos were spaced <10 minutes apart). This gives us confidence that our “event” time limit is not artificially truncating actual event durations by being too short even if the boundary conditions are not absolutely precise.

**Figure 3:**
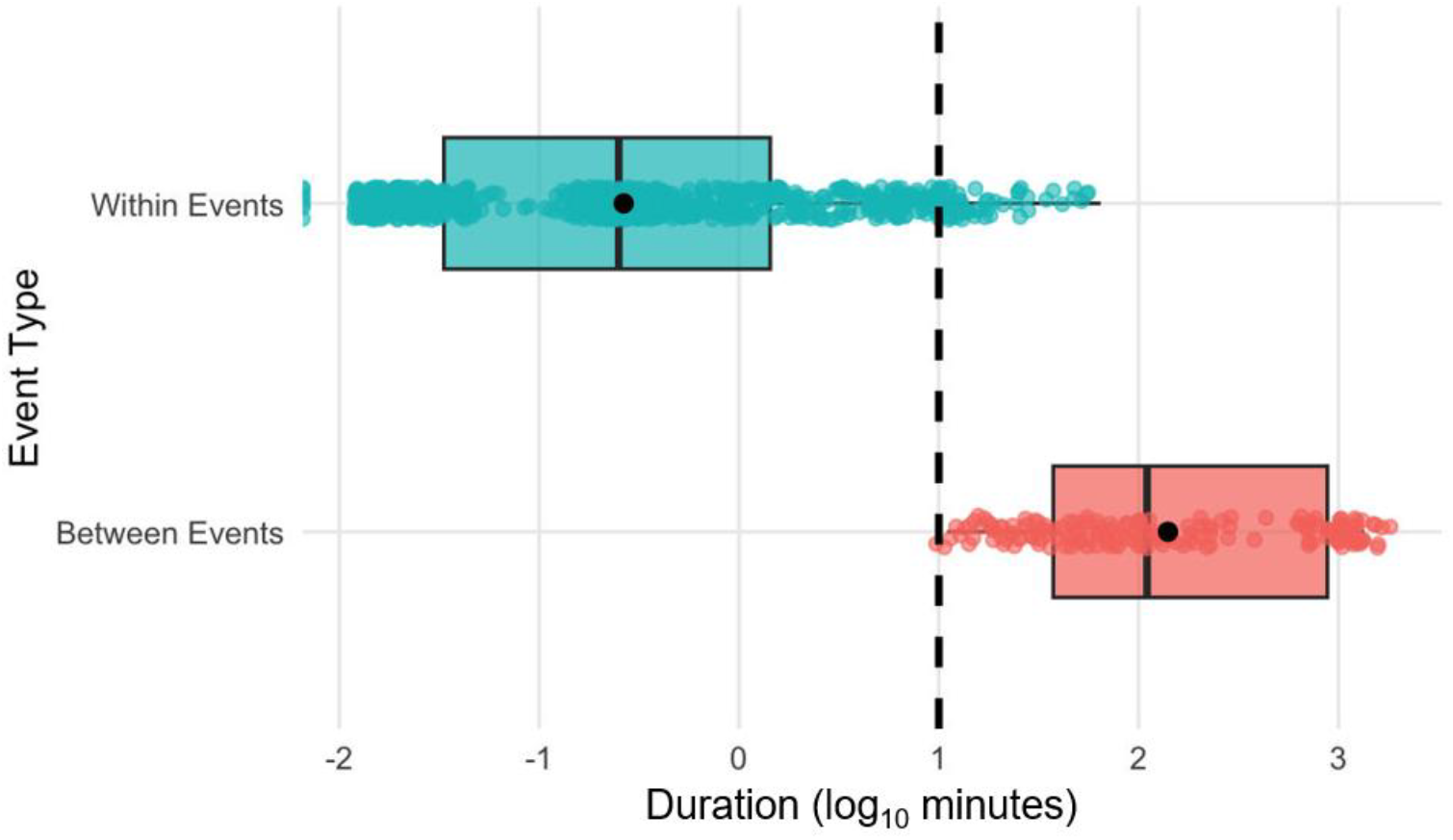
Comparison of event durations vs. interval between events. The number of times (*occurrences*) that within and between event times fall under specific durations (minutes). The dashed line represents the 10-minute threshold. Note that the “within event” duration can be >10 minutes if each consecutive image occurs under this threshold.

### Assessing training success

Volunteers took on average 2.5 hours to get through the initial batch of 50 randomly selected photos but decreased to 30 minutes by the fourth batch. The most used physical attributes to determine sex were genitalia (when seen), back shape, and head bulbosity (Supplemental material, appendix 1). For age, relative size and pigmentation were most used (Supplemental material, appendix 1).

A >90% accuracy rate was reached amongst group 1 when compared to an expert classifications (Böhner et al. 2023). The accuracy of the second set of volunteers (group 2) increased with each sequential batch, going from <70% to 82.6% by the fourth batch (Figure 4). Separate Fleiss’ kappa values were calculated for group type, group size, and event to ensure precise codes from each volunteer (Table 3). Overall agreement among volunteers was lowest for group composition (*K* = 0.247, p<<0.05) and highest for group size (*K* = 0.776. p<<0.05).

**Table 3:**
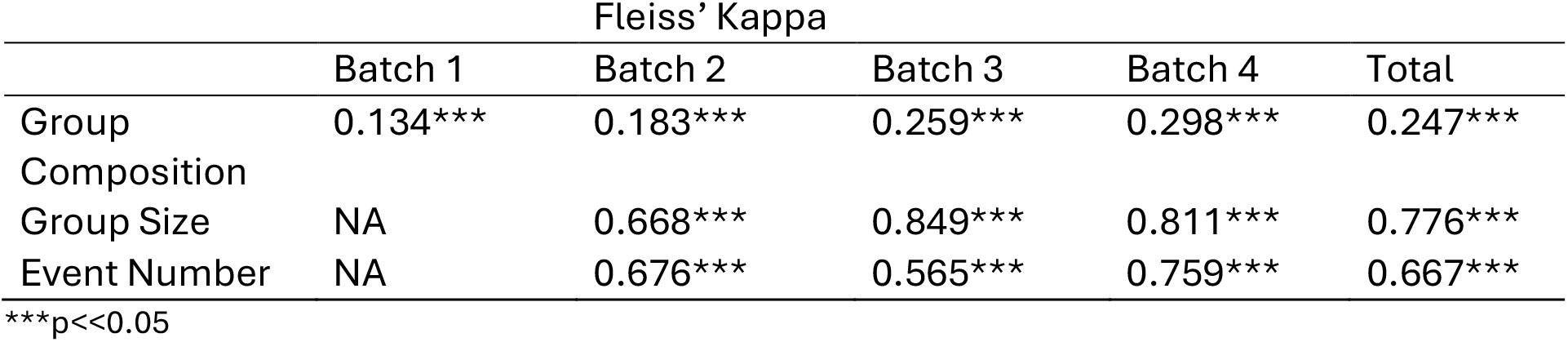
Fleiss’ kappa values for each batch and each category. Group size and event number were not coded for in batch 1.

**Figure 4:**
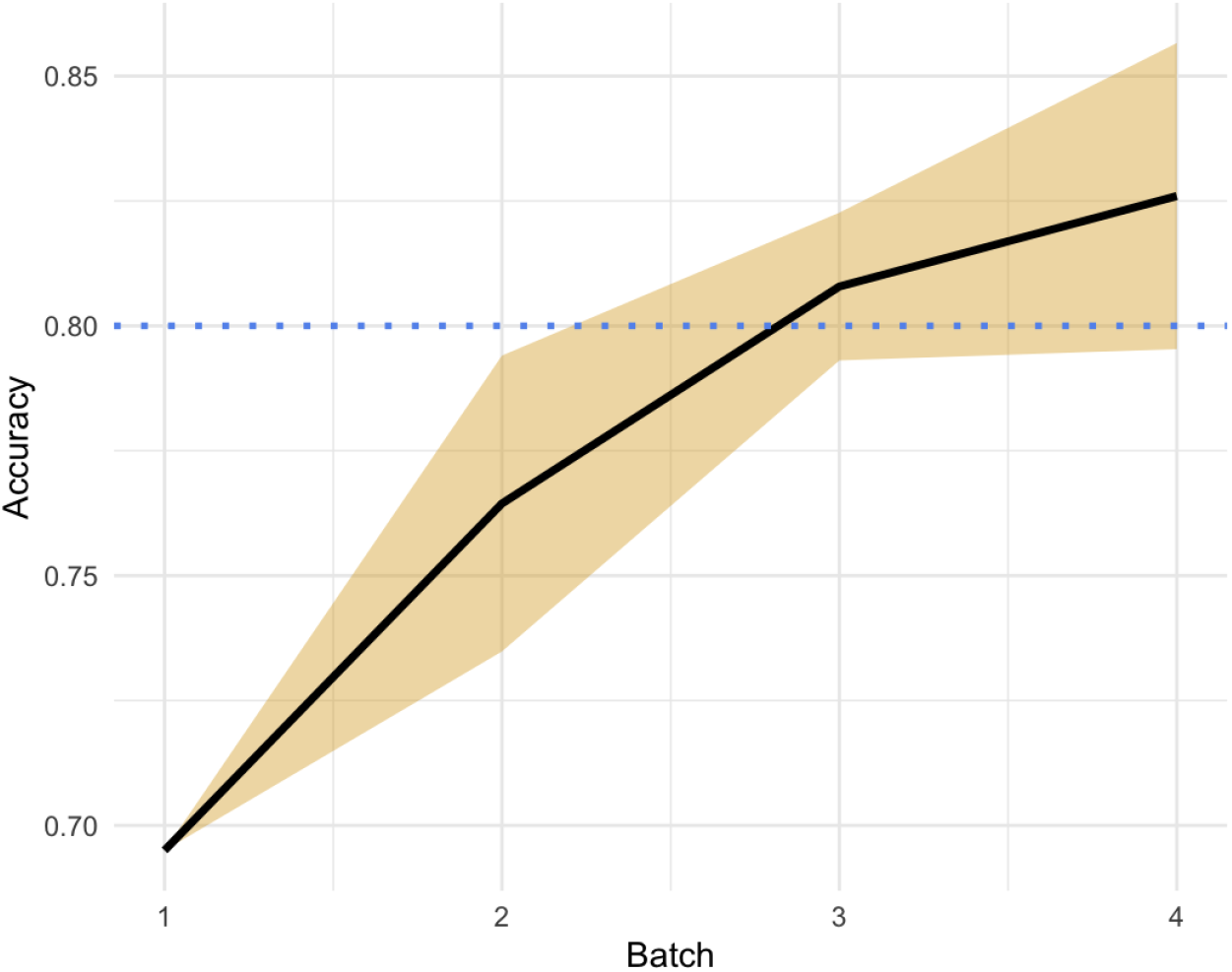
Accuracy across batches. Accuracy over time shows marked improvement among volunteers compared to experienced observers. 80% represents a commonly accepted threshold for “high” accuracy.

## Discussion

The usefulness and popularity of trail cameras in ecological research has grown, generating a large volume of data. While sophisticated automated tools exist for the classification of such images and there has been growing interest in their utility for monitoring species, and elephant in particular (Brickson et al., 2023), they remain largely untested and still face substantial limitations in their utility for making classifications that require interpretation on the basis of multiple contextual cues or refined distinctions among conspecifics. To generate raw training data necessary for these tools (de Silva et al., 2022), and until workflows using them can be validated, there remains a great need for trained human judgement. These labor intensive and limited by the supply of experienced observers. Reliance on untrained individuals suffers from lack of rigor and replicability. To overcome these issues, this study offers a systematic protocol for developing training materials by working with volunteers who have no prior experience. It proceeds in 3 stages:1) initial development of the classification scheme by experienced observers, 2) preliminary training of volunteers and development of a guidance document 3) secondary training and testing of volunteers, together with refinement of the guidance.

Locations and study contexts may vary in the criteria necessary for distinguish among putatively independent events that are captured by cameras. These criteria should be based on knowledge of the taxa and local conditions, validated with test sample. In this study, we distinguished events on the basis of species identity and temporal structure. The vast majority (90.2%) of elephant event durations were well within the threshold of 10 minutes, with the median interval between events (110.22 minutes) being approximately 440 times greater than the median event duration (0.25 minutes). We could use a relatively short 10-minute threshold because prior research on this population indicated elephants occur at very high densities and move together in closely-spaced groups (de Silva et al., 2011a; de Silva et al., 2011b). Even so, it is possible that some ‘solitary’ individuals were in fact socially associated with individuals who passed by >10min before or after them. Other species such as African forest elephants move maintain inter-individual distances of hundreds of meters, and may not move in groups, thus event thresholds might be lower (Beirne et al., 2021). Conversely, if the target species exhibits spatiotemporal coordination/avoidance on larger scales (e.g. many carnivores; dispersed individuals who are in vocal contact or are known to be agonistic/territorial with conspecifics), the event threshold may need to be higher. Depending on the species and area of focus, the event definition can be tested and modified systematically using this approach.

Our training process achieved surprisingly high levels of accuracy and precision among volunteers who had had no prior experience observing elephants, and they did so relatively quickly. Indicators of learning were (a) reduction in time taken for task completion (from 2.5 hours in batch 1 to 30 minutes by batch 4) and (b) improvements in both accuracy and precision across batches. For training artificial intelligence programs, studies like Böhner et al. (2023) suggest 90% as the threshold for proper agreement.

Volunteers in this study achieved over 80% accuracy relative to experts, and the upward trend suggests they likely would have been able to achieve over 90% with additional exposure. A Fleiss’ kappa of 0.298 for the final test of group composition classifications among the six volunteers indicates that there was fair agreement between them, but given this is a relatively small sample size and we recommend training and testing a large set of individuals before application. Fleiss’ kappa values measure the exact agreement between coders. When choosing if an event consisted of a solitary elephant versus group event, the Fleiss’ kappa value was 0.867 by the final batch (almost perfect agreement). Group size and event number had overall kappa values of 0.776 and 0.667, meaning that throughout, observers had substantial agreement on both of these measures (Table 2). Practice enhances performance, therefore we recommend doing 2-3 test batches of camera trap photos using a similar protocol to train any volunteers for coding camera trap photos, regardless of the species. One noteworthy observation is that batch 1, which contained randomly selected images rather than intact image sequences from specific events, were extremely difficult to classify. This highlights the critical utility of contextual information (i.e. comparisons among multiple images within a sequence) for making judgements about group size and composition. Sequential images (or alternately, videos) enhance classification.

We did not attempt to train volunteers to conduct individual identification in this study. Elephants are individually identifiable, primarily using attributes of the ears (de Silva, 2014). Although individual identification from high-quality photographs has been successfully automated, such tools have limited access to training data and have not yet been applied to the lower quality images obtained with camera traps (de Silva et al., 2022). Many images are nevertheless of sufficient quality to identify individuals. Based on experience, individual identification of elephants is a skill that requires long-term exposure and practice to develop even among human observers.

## Conclusion

While the use of trail cameras for observing wildlife has increased substantially, it is important to determine how they are best used for particular species and study contexts. Moreover, while technological improvements allow camera traps to capture thousands of photos per day, AI techniques do not obviate the need for human judgements that make use of nuanced contextual information. These judgements may not be consistent with one another (Zett et al., 2022). These issues can be addressed by developing systematic training protocols tailored to the taxon of interest. These data can be used for a variety of studies, ranging from patch occupancy to population estimation (Ferry et al. 2023). For social species, additional data on group size and composition can provide insight into the demographic health of the population as well as provide a foundation for more detailed studies of behavior, in particular through individual identification as a secondary stage.

## Acknowledgments

We would like to thank Sarah Guidolin, Trevor Ryan, Chaewon Ham, Emma Lam, Jasmine Hubbard, Alison Tran, Dominic Tse, Leisel Geyer, Warren Hui, Ananya Giri, and Gabriel Ulfohn for participating in classifying the trail camera images. We also thank Ravi Kodikara for assisting in data collection. This work was funded by the Association of Zoos and Aquariums and Disney Conservation Grant funds. This work was conducted with permission and in collaboration with the Department of Wildlife Conservation, permit no. WL/3/2/71/17.

## Notes

### Competing Interest Statement

The authors have declared no competing interest.

