## Supplementary methods for "A training protocol for human classification of Asian elephant images from trail cameras"

*Event classification criteria*

Because animals (elephants, cattle) as well as people can move in groups and the same individual could also occur in more than one frame within a set of images, we grouped images into “events”. Events were defined in terms of both species/item type (Table 1) and time intervals. All vehicles were considered independent events. Sequential images containing different species were considered separate events. Different species overlapping in the same frame were also considered independent events and numbered sequentially. Sequential images containing *the same* species were considered distinct if images were separated by a time difference of 10 minutes. This temporal threshold was used because elephants occur in this landscape at high densities (de Silva et al. 2011b) and the spatial proximity criterion used to define social groups in this and other elephant populations typically uses a 500-meter radius to identify individuals that are socially associated (de Silva et al. 2011a, de Silva & Wittemyer 2012). In other words, a 10-minute time difference is sufficient to ensure that all individuals travelling within 500m of one another are counted as being associated with the same event. However, this does not exclude the possibility that some individuals travelling at >10 minutes of one another were in fact part of the same social group.

Elephant groups were further assigned a group size and composition (Table 1). Adults vs. sub-adult age classes were distinguished on the basis of height and physical appearance (de Silva et al. 2011b) but younger age classes were not discriminated for this study. Sex was distinguished by head/back shape and, if visible, genitalia. The protocol document itself is appended below.

#### Appendix 1 – Training Protocol Guideline Document

The following document was created by group 1 volunteers and then used by group 2 volunteers for classifying images unassisted.

### <u>Protocol for Camera Trap Data Entry</u>

#### Differentiating Events

"Events" are used to separate elephants into groups and to record changes in species.

A time difference of over 10 minutes separates different groups. If the same species is seen within 10 minutes of the previous sighting, it should be listed in the same event. Events can last hours, as long as the same species continues to appear within 10-minute intervals without new species showing up.

When multiple species are present in the same frame, it still counts as a single event. There is a second column labeled "second species" to list any frames with multiple species. **Elephants should always take priority and be listed in the first species column when they are present with other species.**

Unknown species are treated as a distinct species. Vehicles and people can also be listed in the second column with other species.

Criteria to start a new event:

- A new species appears in the frame
- Any species leaves the frame
- If the same species reappears after a gap of more than 10 minutes from the last image

Not a new event:

- Empty photos are not considered events and event column is left blank
- Setup photos are labeled as 'Setup'
- If the same species reappears within 10 minutes from the last frame it was seen

#### Group Type

This categorizes the types of "groups" used to classify an event.
It includes the following: *Demographic class*- male, female, mixed group
*Age classes*- adult or subadult. Due to it being difficult to determine age and sex without training, we will be relying on groups of unknown sex and undetermined. Use guides below to help differentiate between the classes.

The classifications are as follows:

AF = Adult female

AM = Adult male

SAF = Subadult female

SAM = Subadult male

FG = Female group

MG = Male group

XG = Mixed group

AUK = Adult unknown sex

SAUK = Subadult unknown sex

UKG = Group of unknown sex

UK = Undetermined

**Musth**

Musth column left blank if no signs of musth are shown. Labeled as Yes or Maybe if signs are present.

Musth is a state of hormonal increase seen in adult male elephants. It is seen as a liquid that flows from the temporal gland above the eyes that can sometimes flow down to the mouth. Extensive urine dribbling, staining the rear legs is also a sign of musth. List as yes if the elephant is clearly in musth. Maybe if you are unsure. Leave blank if the elephant shows no signs of musth.

**Guide to Age**

Females are classified based on reproductive maturity, indicated by having had calves and enlarged mammary glands. Males are classified based on morphology and size because when first reaching reproductive maturity they are socially immature and do not usually mate until later.[3] Additionally, males do not go through any morphological differences after mating.

Aging will be based on the size of morphological female adults. If no full-grown adult female is present causing size/age to be difficult to determine, list as unknown. Only list age if it is clearly identifiable.

Subadult classification is used to categorize all solitary non-adult individuals because juveniles and younger calves are usually seen only with groups.

| Class | Sex | Description | Examples |
|---|---|---|---|
| Adult | Female | Reproductive adult female elephants are ones who have had offspring. Identified through enlarged mammary glands. (10-15 years)[3] Morphologically adults are | Morphological adult: |

| | | | |
|---|---|---|---|
| | | those that have reached maximum height, full-grown adults. (15+ years)[3]<br>Both are equally considered adult females. | 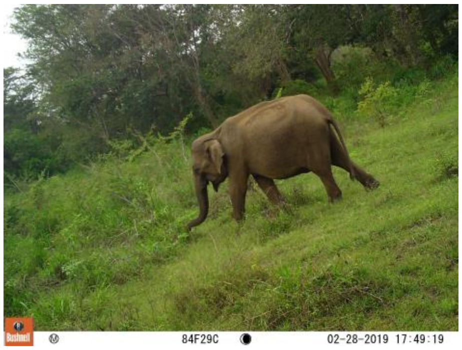<br>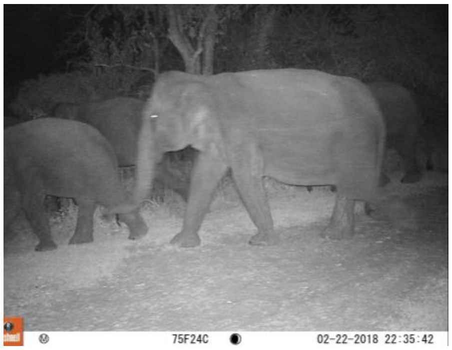<br>Reproductive adult:<br>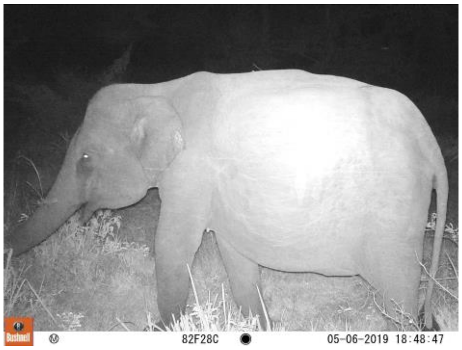<br>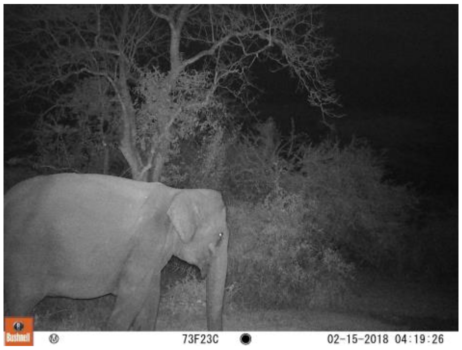 |
| | Male | An adult is any male elephant that is larger than a morphological adult female using the top of the shoulder as the reference point. (15+) | |

| | | | |
|---|---|---|---|
| | | Musth behavior is also an indicator of adult males. | 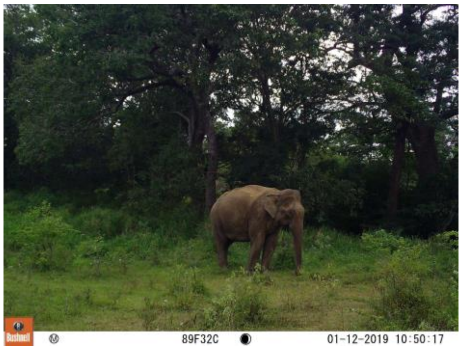  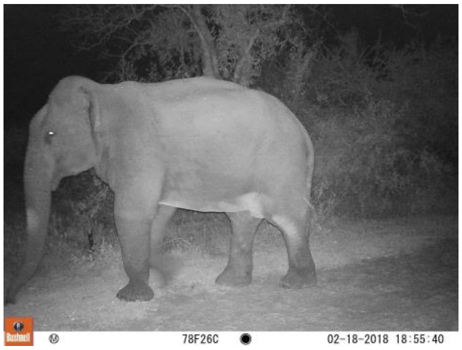  New adult: 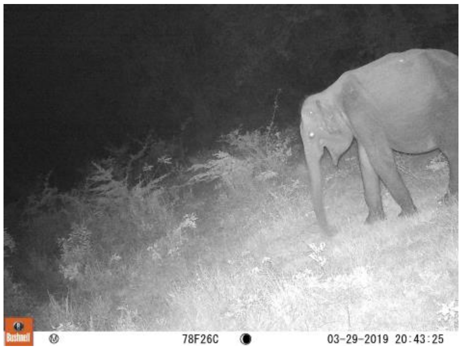  |
| Subadult | Female | Any female who has not had offspring (no enlarged breasts) and is smaller than a morphological adult female. (<10 years) | 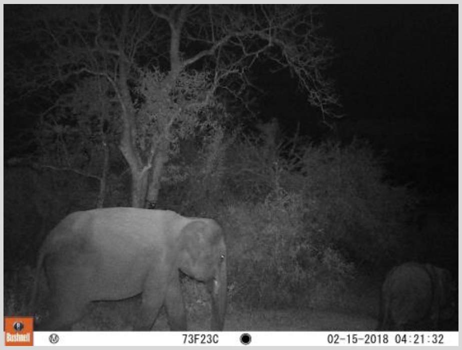  |

| | | | |
|---|---|---|---|
| | Male | Any male the same size or smaller than a full grown adult female, regardless of reproductive maturity. (<15 years) Males reach reproductive maturity at 10 years old but do not become active until later due to social structure.[3] Subadult males are considered reproductive adults but not morphological adults. | 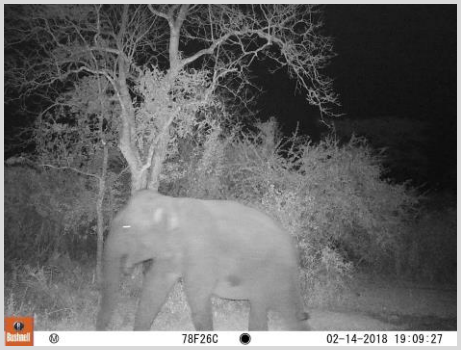  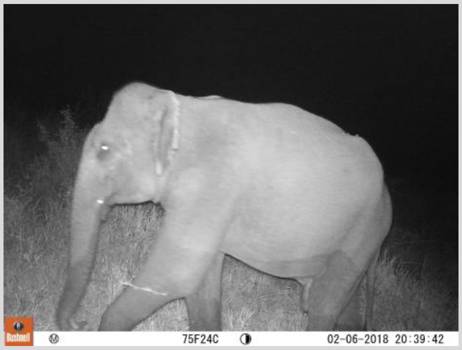  |

Additional Traits:
The following traits are extremely variable and should be used if the above table did not give a clear age classification.

| Trait | Description | Age Class | Example |
|---|---|---|---|
| Depigmentation | Depigmentation on ears and face. Very variable trait and only visible when the elephant is clean. | Adult | 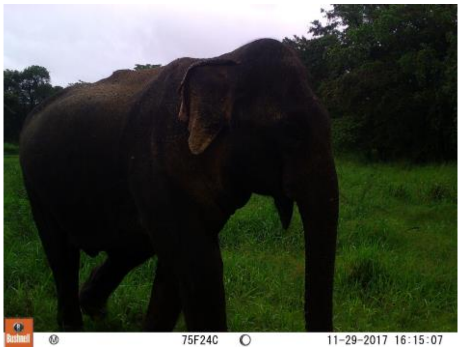  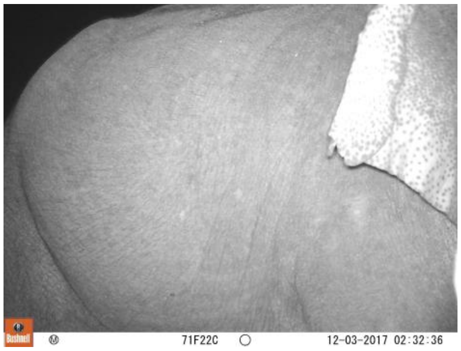  |

| | | | |
|---|---|---|---|
| | Little to no depigmentation | Subadult | |
| Ear Folding | Upper edge of the ears starts to fold inwards after 10 years and progressively gets more prominent | Adult | 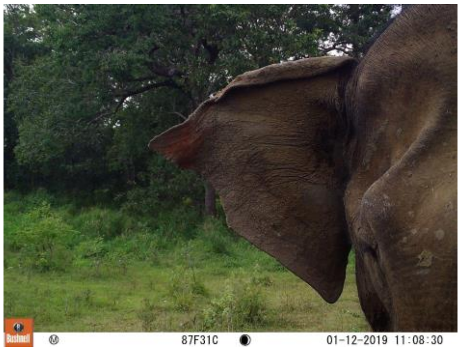  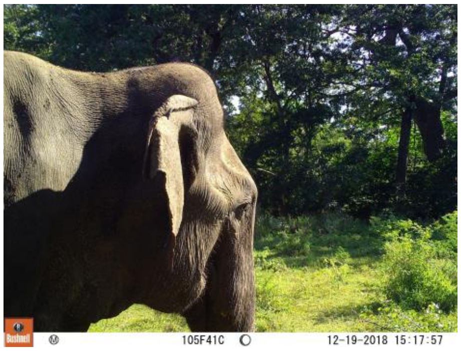  |
| | No ear folding | Subadult | |

Irregular Cases:
There are two dwarf elephants who are adults but that is the stature of a subadult. They are differentiated from the subadults by the body proportion and/or large tusks. These will be coded as adults.

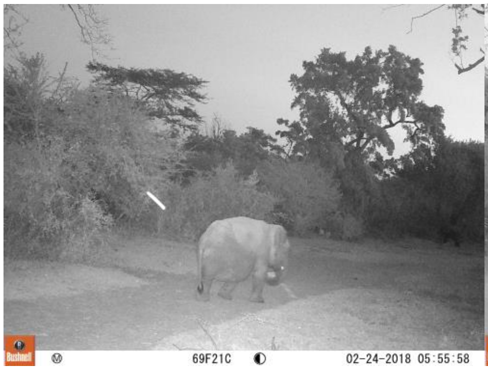

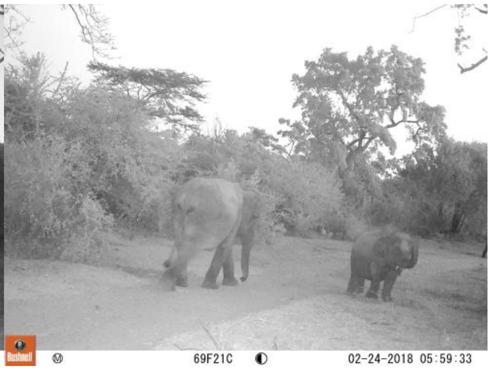


<u>Group pictures for size references:</u>

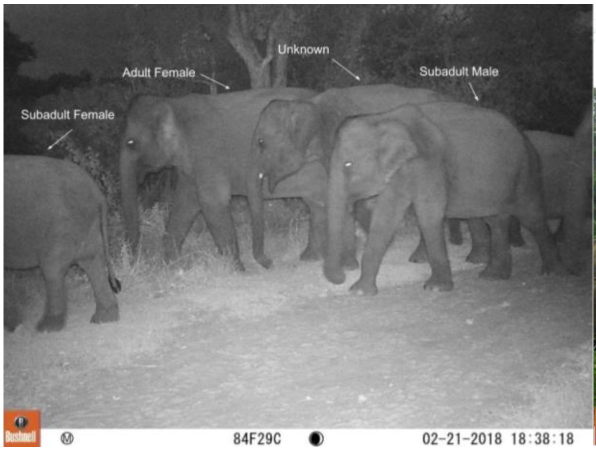

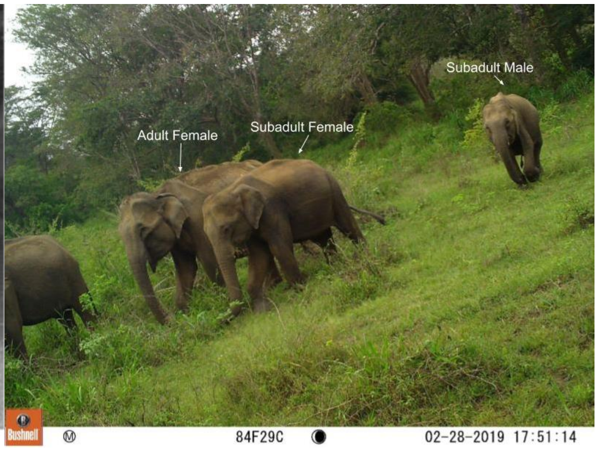


#### Guide to Sexing

Sex classification will be based on multiple features. If sex is not clearly identifiable list as unknown. Make sure you look at all pictures of the elephant before listing a sex, different angles may show different traits and morphology.

Certain traits take priority over others, the traits are listed in descending order of priority. If genitalia, large tusks, mammary glands, or musth is clearly seen then list the sex those traits describe regardless of the other traits.

Ideally the elephant will fit the description for one sex on all traits. This may not always be the case because of individual variation. If the elephant fits the description for both sexes on different traits then it should be listed as unknown.

Ex) Elephant with a bulbous head (male trait) but she has enlarged mammary glands and is seen with a nursing calf. List as an adult female regardless of head shape because mammary glands take priority.

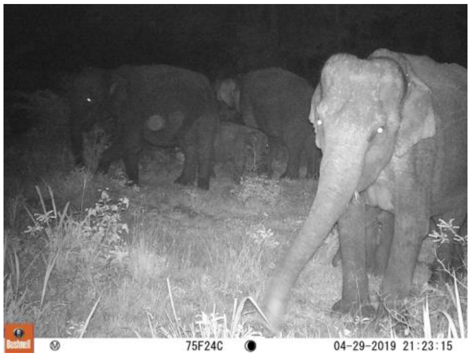

| Traits | Description | Sex | Examples |
|---|---|---|---|
| Genitalia | Obvious penile bulge or visible penis | Male | 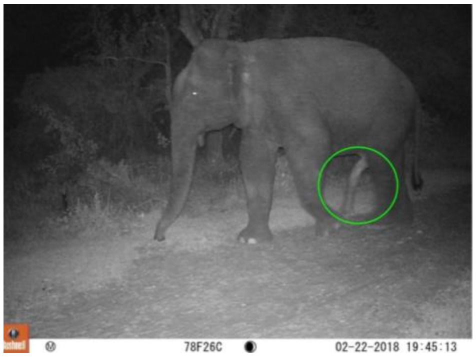<br><br>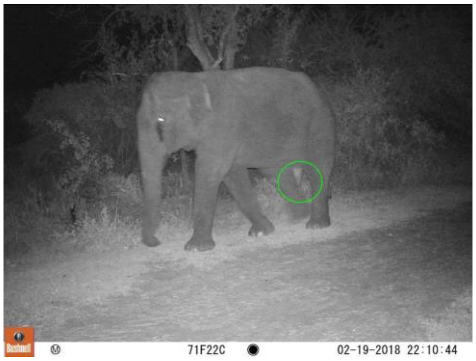<br> |
| | No visible genitalia | Male/Female | Adult males with no visible genitalia:<br>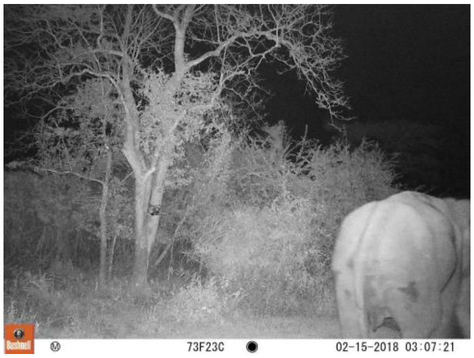<br><br>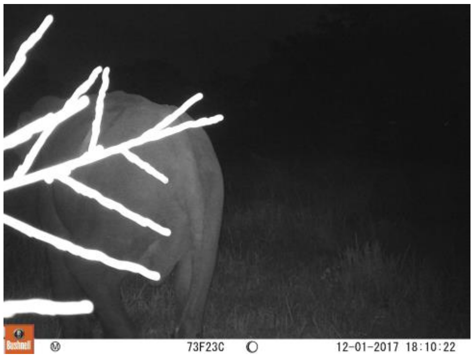<br><br>Adult female: |

| | | | |
|---|---|---|---|
| | | | 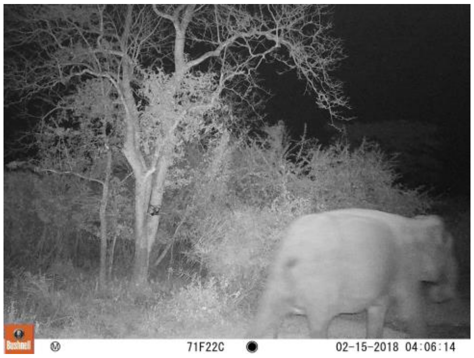 |
| Mammary Glands | No visible mammary glands | Male/Female | Male with no mammary glands:<br>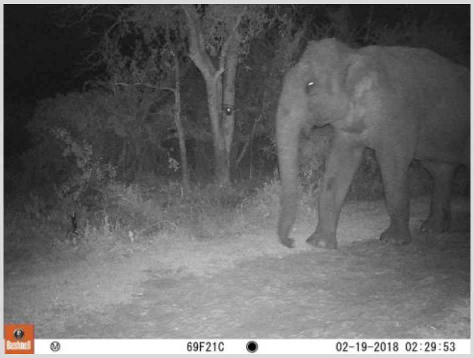<br>Adult female with no enlarged mammary glands:<br>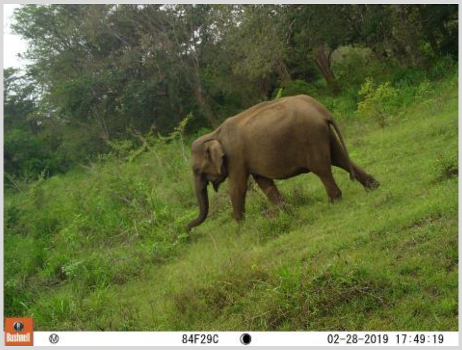 |
| | Enlarge mammary glands. Indicator of adulthood. | Female | 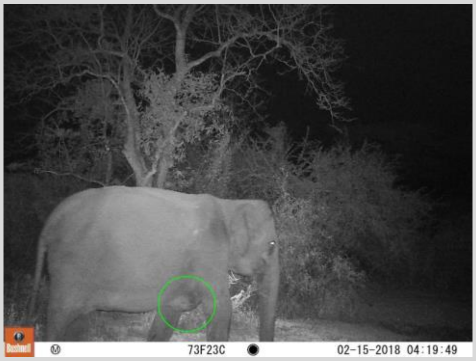 |

| | | | |
|---|---|---|---|
| | | | 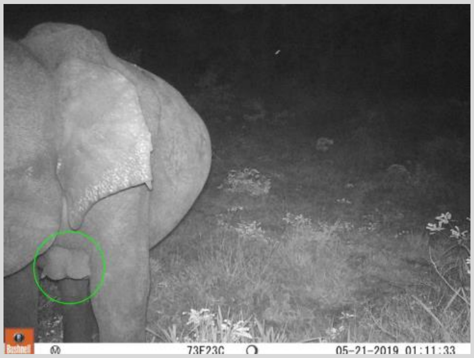 |
| Tusks/Tushes | Large tusks that grow out past the lip. If the lip is not visible but the tusk is visible then the tusk passes the lips and is considered large. | Male | Subadults with tusks:<br>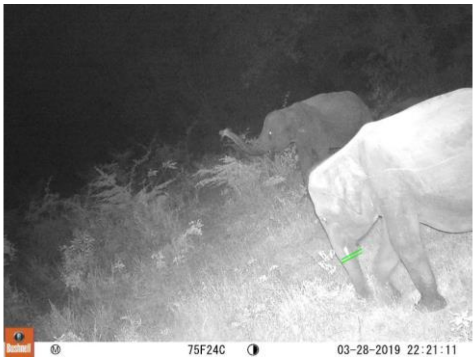<br>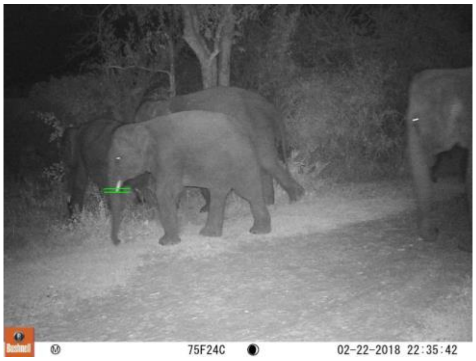<br>Adult with large tusks:<br>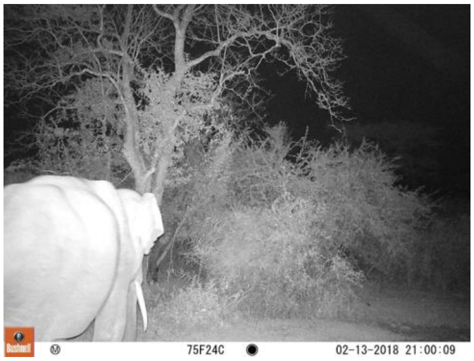 |
| | No visible tusks or tushes that do not pass the lip. | Male/Female | Adult males without tusks: |

| | | | |
|---|---|---|---|
| | | | 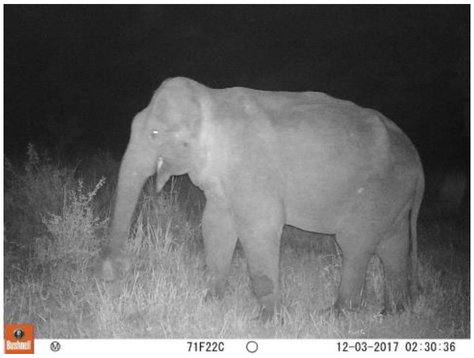<br><br>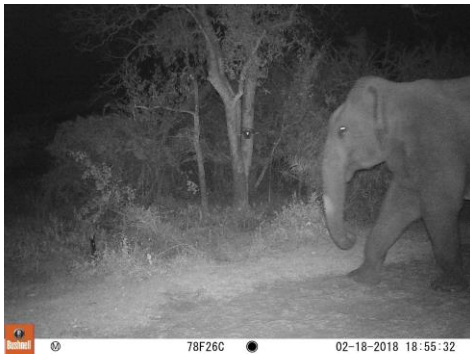<br><br>Male with short tush-like tusk:<br>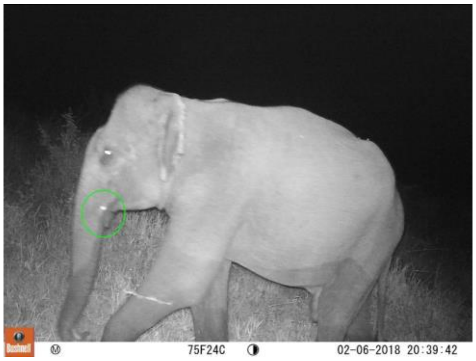<br><br>Females with tushes:<br>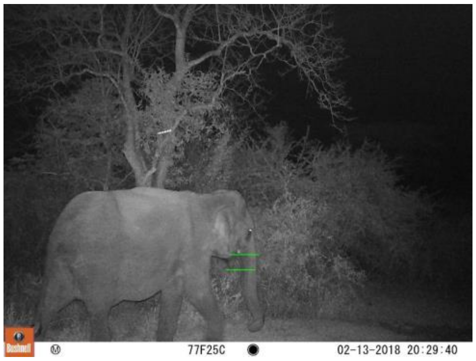<br> |

| | | | |
|---|---|---|---|
| | | | 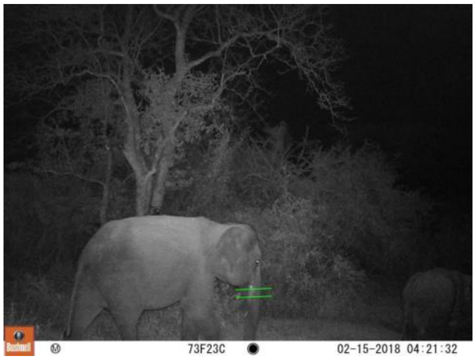  |
| Musth | Extensive temporal and/or urine dribbling. Indicator of adulthood. | Male | 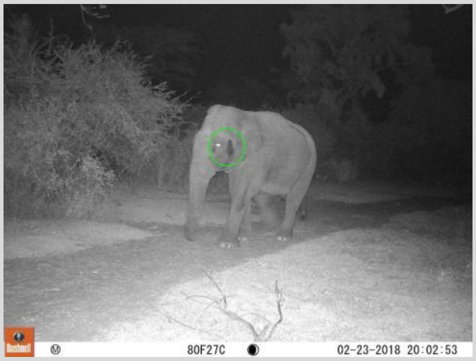  |
| | No noticeable musth. Leave the musth column blank. | Female/Male | Adult male not in musth: 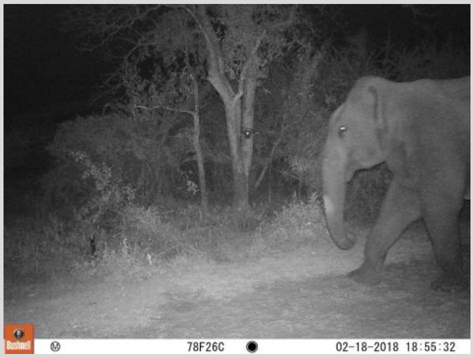  |
| Back Shape | Curved and sloped back. The center of the back is the highest point. Cannot draw a straight line from the end of the back towards the top of the shoulders. | Male | 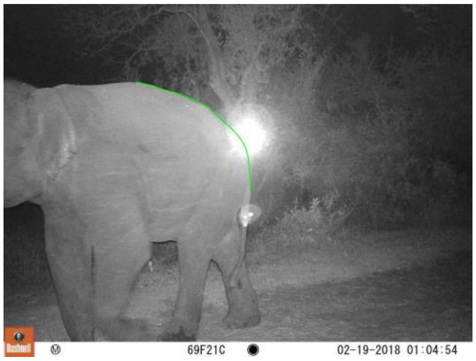  |

| | | | |
|---|---|---|---|
| | | | 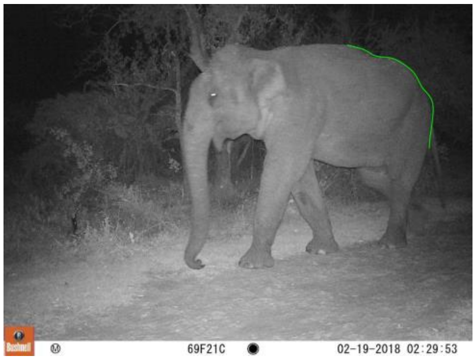 Bushnell 69F21C 02-19-2018 02:29:53 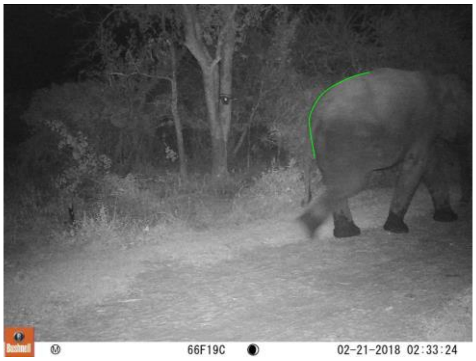 Bushnell 66F19C 02-21-2018 02:33:24 |

| | | | |
|---|---|---|---|
| | Boxy back. A straight line can be drawn from the end of the back to the top of the shoulders. | Female | 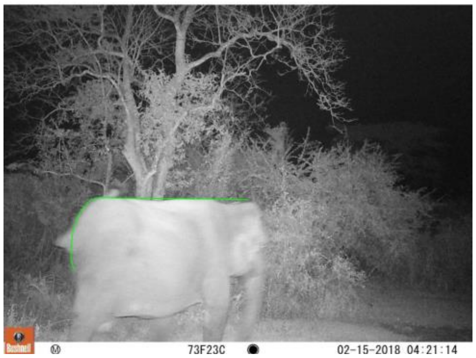 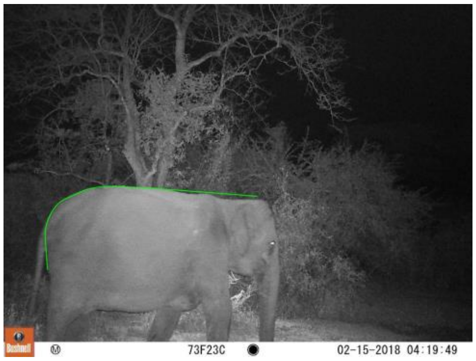 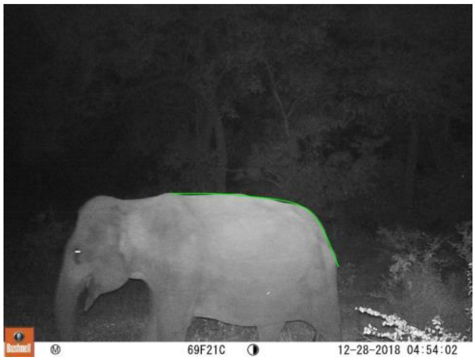 |
| Head Shape | Bulbous head shape caused by prominent parietal domes.<br>Be careful of the angles, different angles can make bulbous heads look flat. | Male | 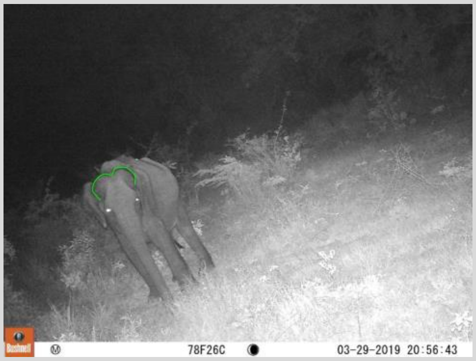 |

| | | | |
|---|---|---|---|
| | | | 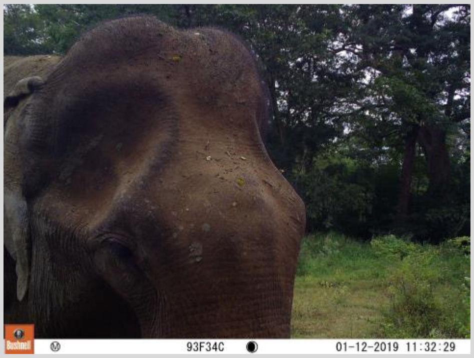  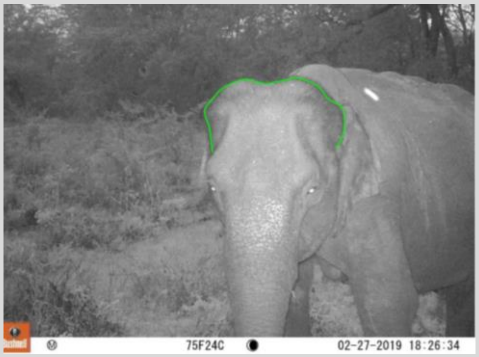  |
| | Boxy head shape with no prominent domes or with small prominent domes.<br>Careful with the angles, different angles can make their heads more bulbous than they are. | Male/Female | 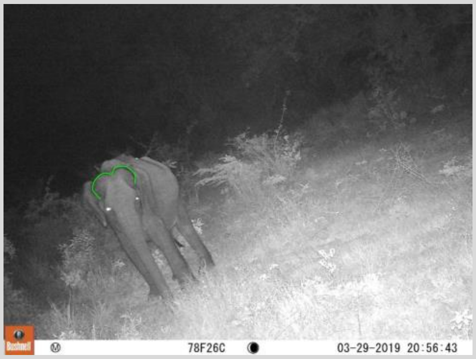  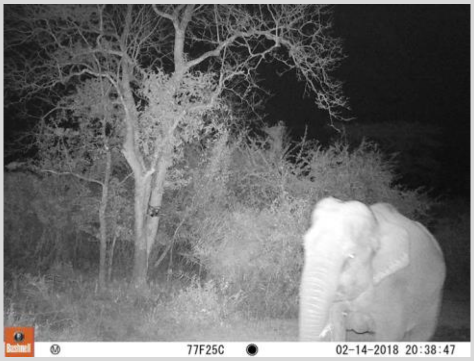  |

| | | | |
|---|---|---|---|
| | | | Male with less prominent domes: 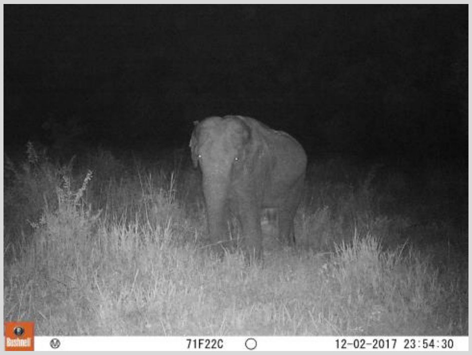  |
| Nasal Protuberance | Nasal point between the eyes comes out in a prominent peak compared to parietal domes. | Male | 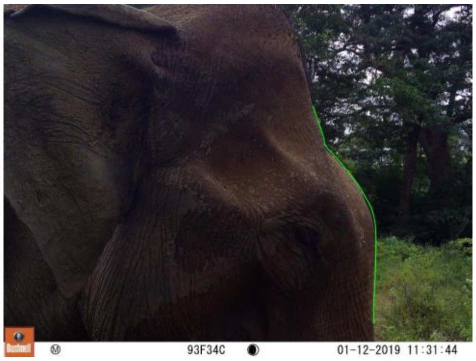  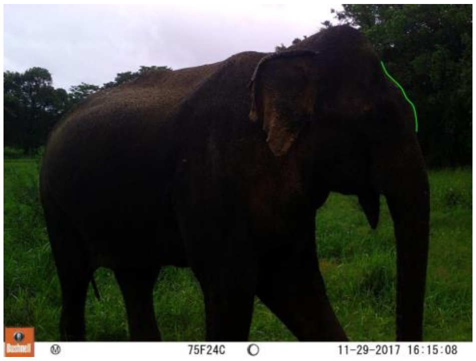  |

| | | | |
|---|---|---|---|
| | No prominent peak. Nasal point is more linear and connected to head domes. | Female/Male | 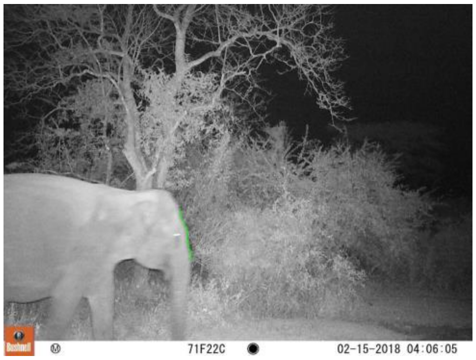  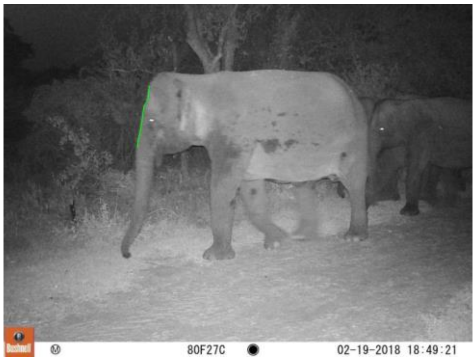  Male without a prominent nasal peak: 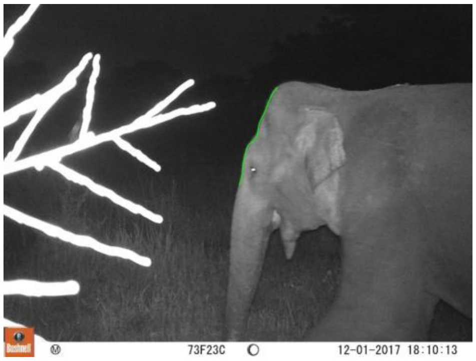  |
| Foreleg to hindleg ratio | Large foreleg to hindleg length difference. Causes the sloped back. | Male | 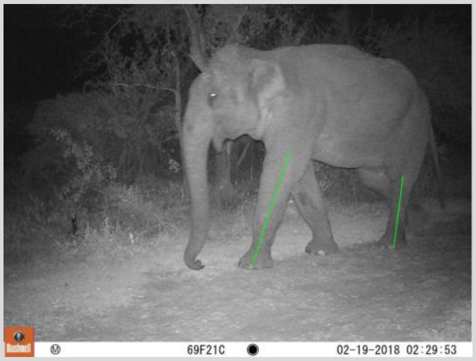  |

| | | | |
|---|---|---|---|
| | Smaller foreleg to hindleg length difference. | Female | 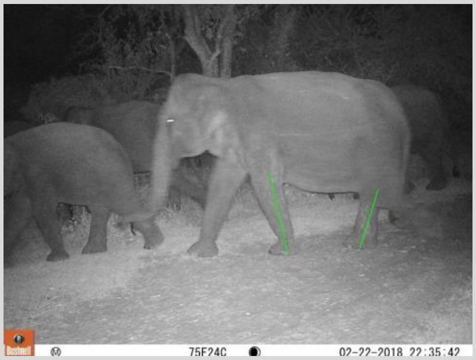 |

#### **Supplemental Images**

Depigmentation:

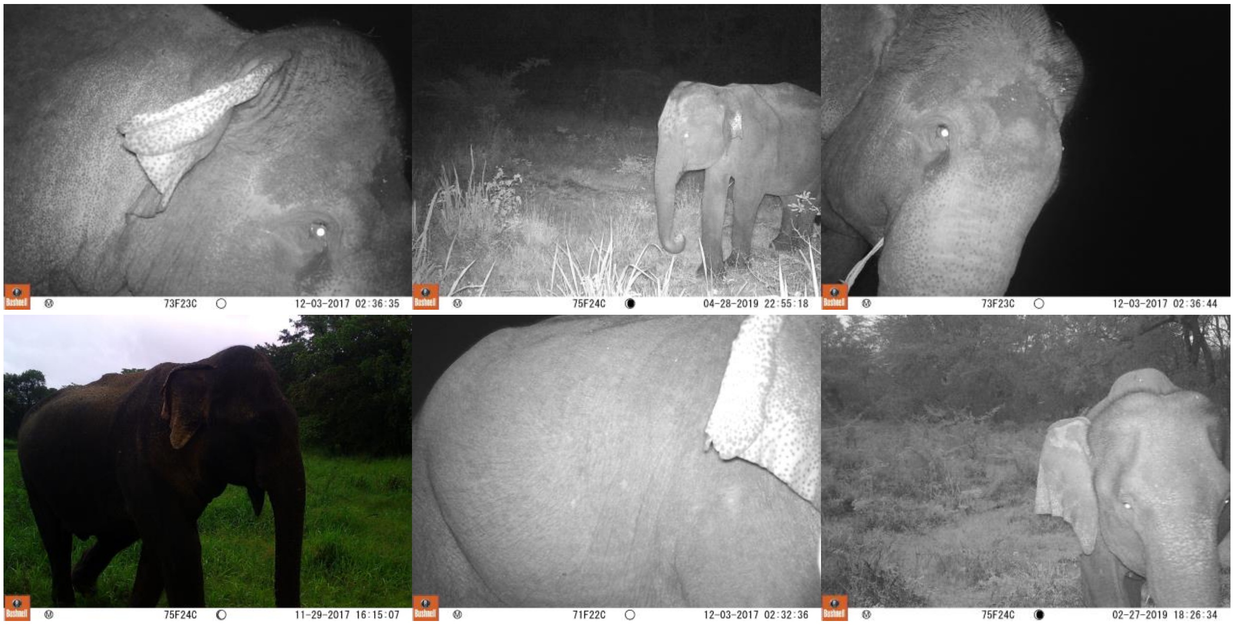


Ear folding:

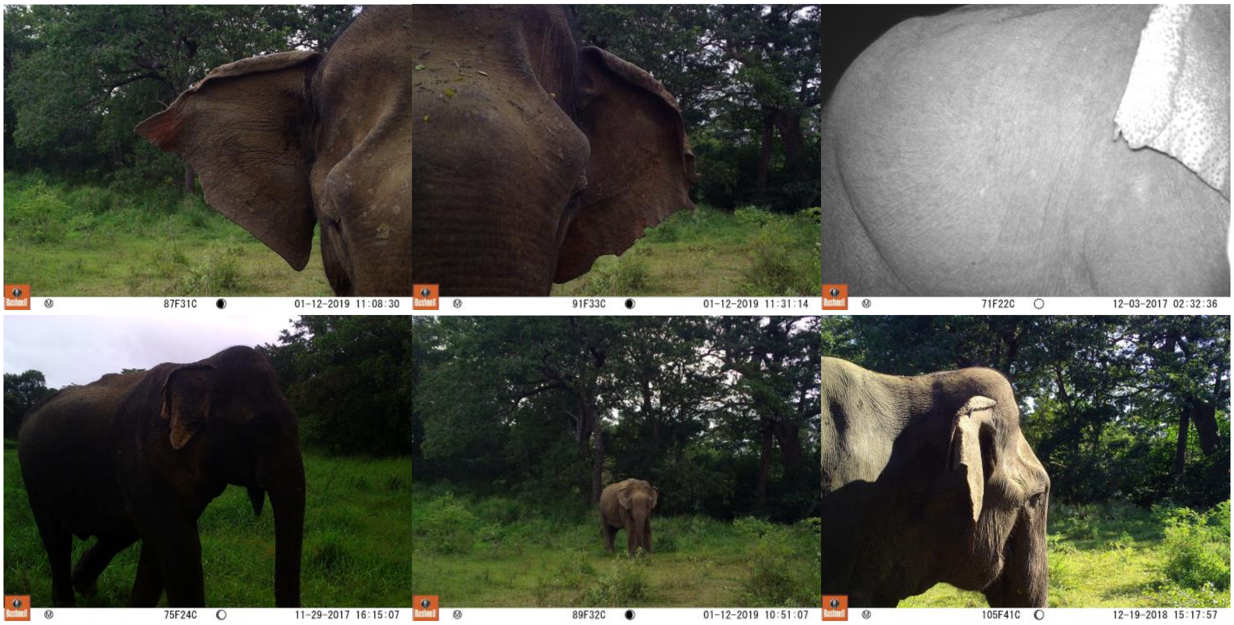

87F31C 01-12-2019 11:08:30
91F33C 01-12-2019 11:31:14
71F22C 12-03-2017 02:32:36
75F24C 11-29-2017 16:15:07
89F32C 01-12-2019 10:51:07
105F41C 12-19-2018 15:17:57


AM:

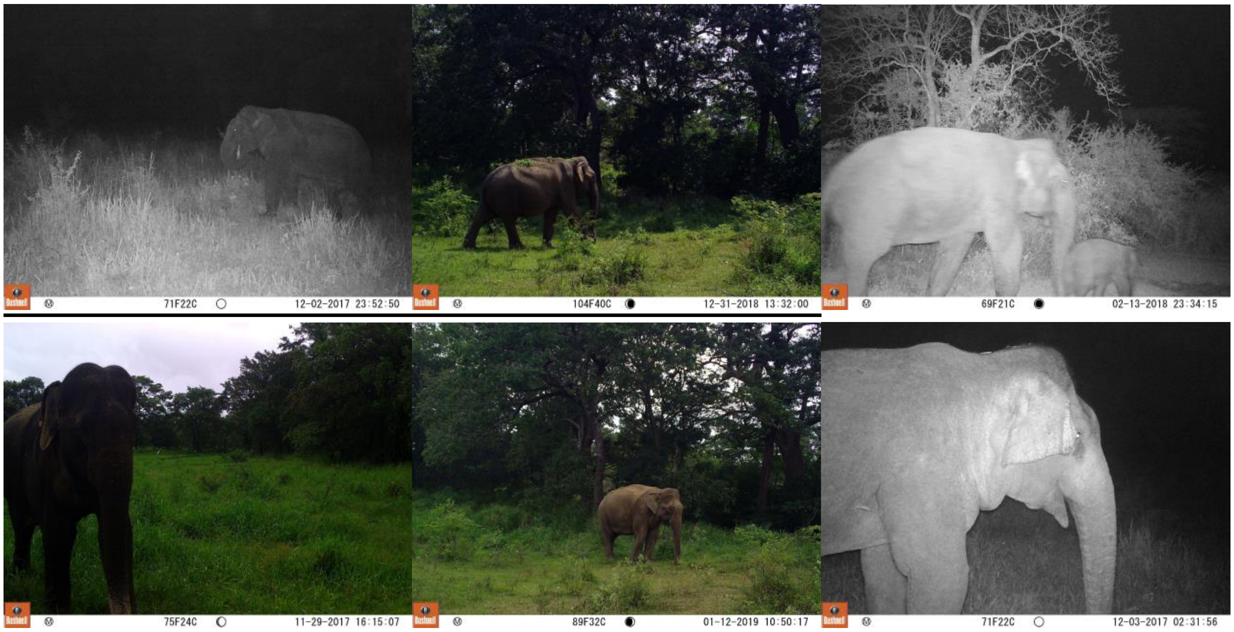

71F22C 12-02-2017 23:52:50
104F40C 12-31-2018 13:32:00
69F21C 02-13-2018 23:34:15
75F24C 11-29-2017 16:15:07
89F32C 01-12-2019 10:50:17
71F22C 12-03-2017 02:31:56


AF:

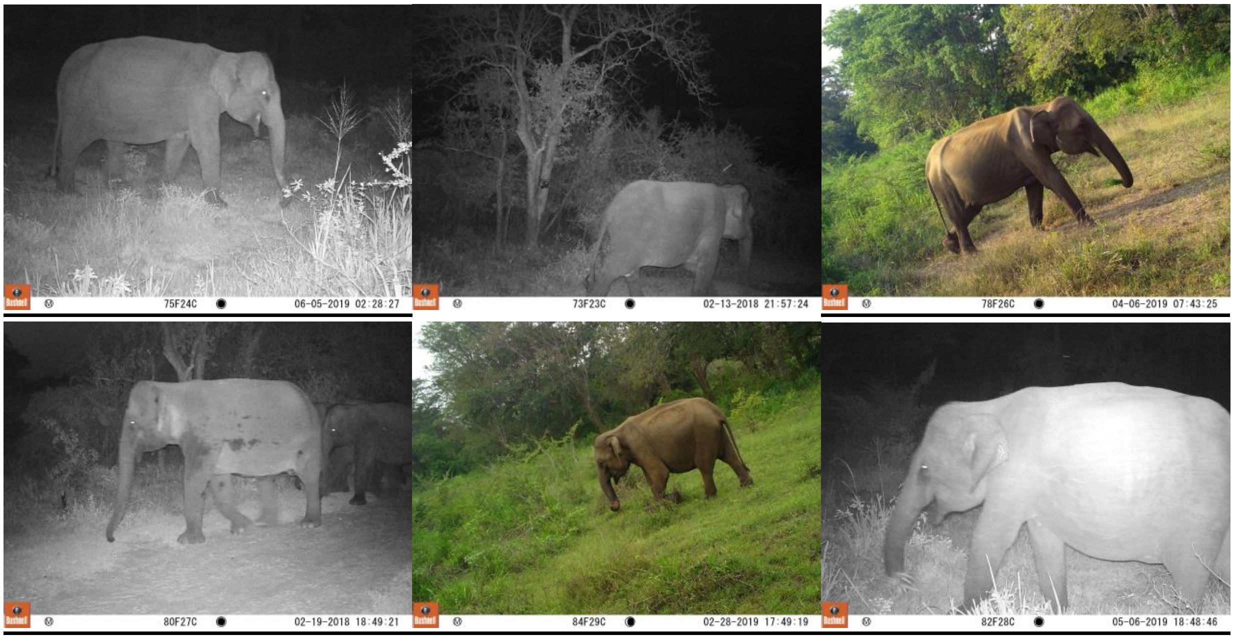


AF with a bulbous head:

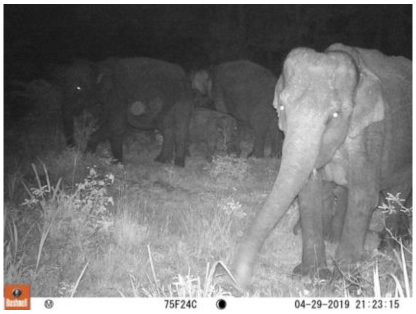


Adult male nasal protuberance from different angles:

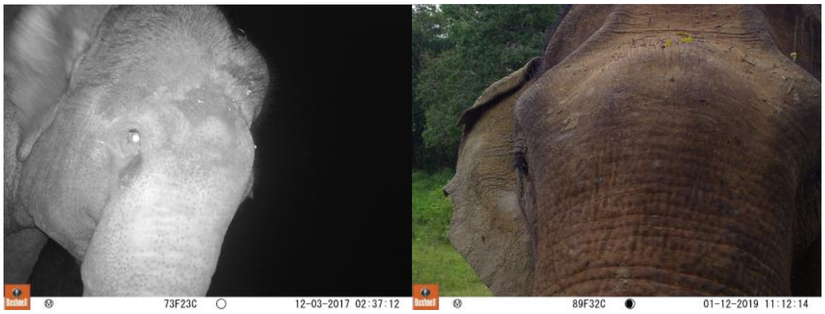


Comparison of head shapes between female and male:

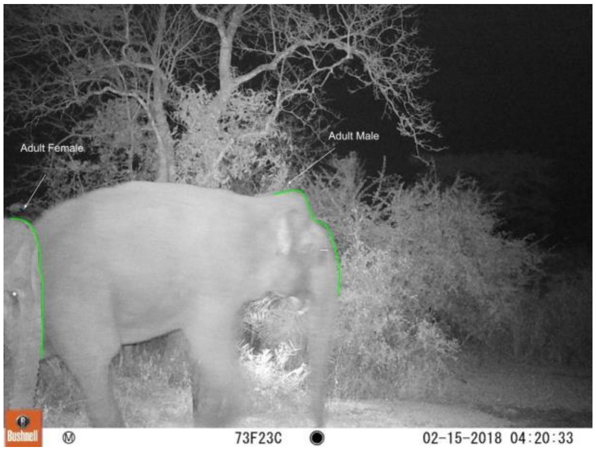


Adult vs subadult males:

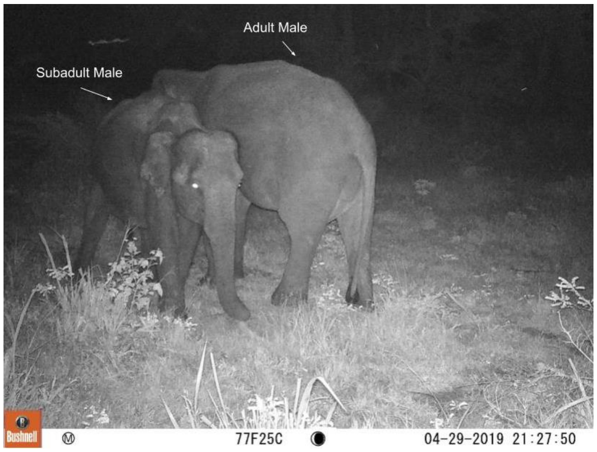
